# The effect of wheat genotype on its microbiome is more evident in roots than rhizosphere and is strongly influenced by time

**DOI:** 10.1101/2022.11.10.515967

**Authors:** Liliana Quiza, Julien Tremblay, Antoine P. Pagé, Charles W. Greer, Curtis J. Pozniak, Rong Li, Brenda Haug, Sean M. Hemmingsen, Marc St-Arnaud, Etienne Yergeau

## Abstract

Crop breeding has traditionally ignored the plant-associated microbial communities. Consideration of the interactions between plant genotype and associated microbiota is of value since different genotypes of the same crop often harbor distinct microbial communities which can influence the plant phenotype. However, recent studies have reported contrasting results, which led us to hypothesize that the effect of genotype is constrained by time (growth stage, year) and space (plant compartment). To test this hypothesis, we sampled bulk soil, rhizosphere soil and roots of 10 wheat genotypes, twice per year, for 4 years. DNA was extracted and regions of the bacterial 16S rRNA and CPN60 genes and the fungal ITS region were amplified and sequenced. The effect of genotype was highly contingent on the time of sampling and on the plant compartment sampled. Only for a few sampling dates, were the microbial communities significantly different across genotypes. The effect of genotype was most often significant for root microbial communities. The three marker genes used provided a highly coherent picture of the effect of genotype. Taken together, our results confirm that microbial communities in the plant environment strongly vary temporally and spatially and that this can mask the effect of genotype.

## Introduction

Crop breeding has traditionally been carried out under high input conditions and at the same time, ignoring plant-associated microbial communities [1–3], leading to the view that modern genotypes have lost their ability to associate with nutrition-beneficial microbial communities when grown under low inputs [4–9]. Since many plant-associated microorganisms can positively affect the plant phenotype by increasing nutrition, deterring pathogens, promoting growth and reducing stress, their absence could significantly hamper efforts toward sustainable agriculture. However, plant-associated microorganisms are in turn influenced by seasonal variation in environmental conditions [10], plant developmental stage [11, 12], plant compartment [11, 13], and their interactions [14], which probably shape the strength of the association between microbial communities and particular plant genotypes.

Microbial communities were shown to vary with plant developmental stage but also with environmental conditions throughout the growing season. In the field, both sources of variation are confounded, making it difficult to tease apart the influence of development stages and plant and microbial responses to variation in environmental conditions. However, studies under controlled environmental conditions showed that plant developmental stage does influence the microbial community diversity [11, 15, 16] and activity [12], through changes in the composition of the root exudates [17, 18]. Similarly, field studies have shown that soil microbial communities are influenced by time through changes in environmental conditions, such as precipitation patterns [19] nutrient availability [20], and temperature [21]. For instance, Wang et al. [22] showed that the effects on microbial communities of dry spells followed by rewetting, dwarfed the effects of reduced soil water content and of different wheat genotypes. Plants also respond to environmental conditions by modulating their rhizodeposition [23, 24], providing an indirect way for seasonal variation in environmental conditions to affect microbial communities.

Plant compartment is often reported as the most dominant factor influencing the diversity of plant-associated microbial communities [11, 25, 26]. One of the most reported patterns is the difference between the rhizosphere and bulk soil microbial communities, also dubbed the rhizosphere effect [6, 27–29]. Likewise, root, rhizosphere and aboveground plant microbial communities are distinct [11, 30, 31], these latter sharing similarities with the seed microbial communities [32]. These differences are due to a combined selective pressure of quantity and quality of nutrients [33], microbial capacity to invade plant tissues, and plant immune response to invaders [28, 34, 35]. For instance, assembly in the rhizosphere is linked to the presence of various chemicals exuded by the roots [18, 36, 37], to the capacity of the microbes to consume these exudates or react to them and to form biofilms [38, 39]. Invasion of the plant tissue, such as the root endosphere, requires the microbes to evade plant defenses and to adapt to life inside tissue where nutrient sources are highly unbalanced.

A recent study from our group showed that spatial and temporal factors interact to modulate microbial communities [11]. This suggested that depending on the plant compartment, the effect of plant developmental stages on the microbial communities is not identical. It would therefore be expected that an effect such as plant genotype, that is generally reported to have a more subtle effect on microbial communities than environment or plant compartments [19, 40], would be influenced and even constrained by these factors. One of our recent studies showed that the rhizosphere metagenomes of 10 different wheat genotypes were nearly identical [9]. However, previous work from our group did highlight significant differences between wheat genotypes in term of function, community structure and composition for wheat growing in pots in a growth chamber [40, 41], in commercial fields [42], or in an experimental field [19]. We therefore hypothesized that the effect of wheat genotype might show some spatio-temporal variability, being only visible at certain growth stages, in certain plant compartments or under certain environmental conditions, which could explain the lack of significance observed in Quiza et al. [9]. Here, we expanded the analysis presented in Quiza et al. [9] by sampling the same field experiment, but twice per season, over four growing seasons, and by including root and bulk soil samples in addition to the rhizosphere soil. We used amplicon sequencing targeting the bacteria and archaea (16S rRNA and CPN60 genes) and the fungi (ITS region) to characterize the entire soil microbial community. Our study sheds light on the interactive effects between plant genotype, time and space and suggests that the genotype effect is plant compartment and time dependent.

## Materials and methods

### Experimental design

A field experiment was conducted from 2013 to 2016 at the Nassar Crop Research Farm of the University of Saskatchewan, Saskatoon, Canada, that has been managed for more than 50 years to conduct experiments under low fertilization conditions. We selected 10 wheat genotypes that were introduced over the period 1845 through 2009. These included six *Triticum aestivum* or bread wheat genotypes of the Canada Western Red Spring (CWRS) class (Red Fife (introduced in 1845), Marquis (1911), CDC Teal (1991), AC Barrie (1994), Lillian (2003), CDC Kernen (2009)) and four *T. turgidum* ssp. *durum* or durum wheat genotypes of the Canada Western Amber Durum (CWAD) (CDC Stanley (2009), Pelissier (1929), Strongfield (2004), and CDC Verona (2008)) (https://grainscanada.gc.ca/en/grain-quality/grain-grading/wheat-classes.html). The experiment was arranged in three randomized blocks, each consisting of ten 6.2 m^2^ plots to which the cultivars were randomly assigned each year. Each plot contained eight rows spaced at intervals of 20 cm and were seeded at 320 seeds m^−2^ on May 25, 2013, June 2, 2014, May 20, 2015, and June 7, 2016. To minimize the effect of the seed source on plant performance, all cultivars were grown from seeds harvested from a common field under low fertilization. To minimize any erroneous effect on productivity measurements due to poor seedling establishment, 15 kg ha^−1^ of 11-55-0 (NPK) fertilizer was added at seeding. Weather conditions (average daily temperature and total monthly precipitation) for Saskatoon (World Meteorological Organization station ID: 71496, 52°10’25.000” N, 106°43’08.001” W) for the duration of the experiment (summer 2013-2016) are presented in Table 1.

**Table 1.** Average daily temperature and total precipitation for June-September in Saskatoon, years 2013-2016

|  |  | Avg. daily<br>temperature | Monthly<br>precipitation<br>sum |
| --- | --- | --- | --- |
| 2013 | June | 20.9 | 117.7 |
|  | July | 23.6 | 35.6 |
|  | August | 26.4 | 14.9 |
|  | September | 22.9 | 15.4 |
| 2014 | June | 19.4 | 94.8 |
|  | July | 24.5 | 44.5 |
|  | August | 24.6 | 18.5 |
|  | September | 19.4 | 10.7 |
| 2015 | June | 24.8 | 13.6 |
|  | July | 26.7 | 84.3 |
|  | August | 24.1 | 45.2 |
|  | September | 18.5 | 50 |
| 2016 | June | 24.7 | 49.7 |
|  | July | 25.3 | 58.6 |
|  | August | 23.3 | 70.2 |
|  | September | 19.2 | 24.1 |

### Sampling

Samples from the bulk soil, rhizosphere and root were collected twice per year during four consecutive years (2013-2016), on July 2, 2013, August 26, 2013, July 7, 2014, September 3, 2014, June 22, 2015, August 17, 2015, July 5, 2016 and August 22, 2016. This corresponded approximately to the stem elongation (June and July) and dough development (August and September) growth stages. Three soil cores containing five or more plants were collected in each plot and refrigerated on the sampling day. The soil cores were pooled before processing on a clean piece of bench cover. The shoots were cut with sterilized scissors 1 cm above the soil line. The roots were gently separated from the bulk soil by removing all the lumps and leaving the roots with very little soil adhering to them. The roots were cut off from the remaining stem and crown of the plant with sterilized scissors; they were immediately transferred, with the adhering soil, to a 500 ml Erlenmeyer flask containing 200 mL of sterile PBS buffer. All the roots from one plot were placed in one flask. The flask was placed on a shaker at 150 rpm, at 22°C for 25 min. The roots were removed from the PBS solution and rinsed with distilled water until completely clear, finishing the cleaning with a final rinse of sterile water. The PBS containing the remaining soil from the roots was then centrifuged at 2000 ×g for 5 minutes, after which the supernatant was discarded, and the rhizosphere soil collected. The remaining bulk soil was fragmented into pea-size pieces, mixed and subsampled in 50 ml conical tubes. The roots, rhizosphere soil and bulk soil were immediately stored at −80°C until extraction.

### DNA extraction

Total DNA was extracted from 250 mg of bulk or rhizosphere soil with MoBio’s PowerSoil DNA Isolation Kit (MO BIO Laboratories, Inc, Carlsbad, CA). Three grams of root material were pulverized with the Geno Grinder (HORIBA Canada Inc., London, Ontario) and 50 mg were extracted using the PowerPlant Pro DNA Isolation Kit (MO BIO Laboratories, Inc.). DNA samples were quantified by fluorescence detection using the Life Technologies Qubit® dsDNA HS quantitation kit (Invitrogen, Waltham, MA). Libraries for sequencing were prepared according to Illumina’s “16S Metagenomic Sequencing Library Preparation” guide (Part#15044223Rev.B). Amplicon libraries for the bacterial 16S rRNA gene were prepared using primers F343 and R803 [43] whereas for the fungal ITS1 region the primers ITS1F and 58A2R [44] were used. The cpn60 UT was amplified using the type I chaperonin universal primer cocktail containing a 1:3 ratio of H279/H280:H1612/H1613 as previously described [45, 46]. The three pools were then loaded on an Illumina MiSeq sequencer and sequenced in-house using a 600-cycles MiSeq Reagent Kit v3.

### Bioinformatics and statistical analyses

Sequencing data were analysed using Amplicon Tagger [47] as described previously in [48]. All statistical analyses and figure generation were performed in R v.4.0.2. The similarity between samples due to the ASV relative abundance was visualized by principal coordinate analysis (“cmdscale” function) based on Bray-Curtis dissimilarity matrices (“vegdist” function of the “vegan” package) [49]. The effects of genotype, year, growth stage and compartment on the community composition was tested by permutational multivariate analysis of variance (PERMANOVA) using the “adonis2” function of the “vegan” package. For all the significant (P<0.05) and marginally significant (P<0.10) year/compartments/growth stage communities, pairwise comparisons of the cultivars was performed (function “pairwiseAdonis”) [50]. ANOVA analyses were performed with the function “aov” followed by post hoc Tukey’s honestly significant difference (HSD) tests (“agricolae” package) to detect differences in relative abundance of phyla and classes between genotypes through the years.

## Results

### Microbial community structure

The PCoA ordination based on bacterial 16S rRNA gene amplicon ASVs revealed the overriding effect of compartment on the bacterial community structure, with bulk and rhizosphere samples clustering away from roots samples (Fig. 1). Within each compartment, a clear effect of time was visible, with clustering of samples according to sampling year and growth stage, differentiating the 2013-2014 samples from the 2015-2016 samples (Fig. 1). The Permanova confirmed this visual interpretation, with Compartment having the strongest effect on the community structure (highest pseudo-F ratio) followed by Year and Growth stage, and finally by Genotype (Table 2). All these main effect terms were having a significant effect on the community structure, together with many of the interaction terms, some of them including Genotype (Genotype:Year and Genotype:Compartment) (Table 2).

**Figure 1.**
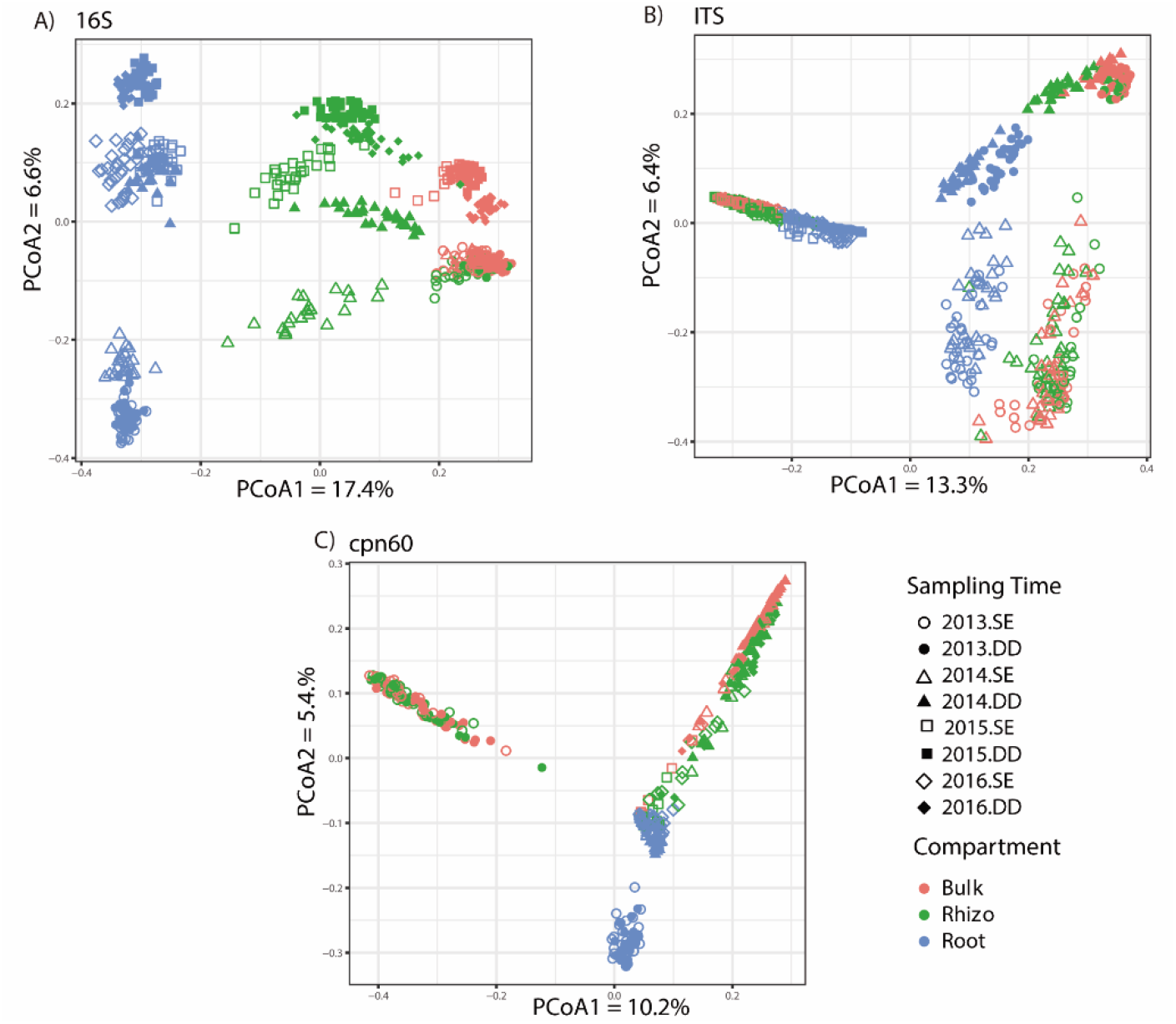
Principal coordinate analysis based on Bray-Curtis dissimilarity between ASV composition derived from A) 16S rRNA gene amplicons, B) ITS region amplicons and C) cpn60 gene amplicons.

**Table 2.** Permanova analysis for the effect of genotype per year, compartment, and growth stage for 16S, ITS and cpn60 amplicon datasets generated from bulk soil, wheat rhizosphere soil and wheat roots collected from 2013-2016 at the stem elongation and dough development stages.

|  | *16S* |  | *ITS* |  | *cpn60* |  |
| --- | --- | --- | --- | --- | --- | --- |
|  | *F* | *P* | *F* | *P* | *F* | *P* |
| Genotype (G) | 1.66 | 0.003** | 1.17 | 0.085· | 1.50 | 0.001 *** |
| Year (Y) | 146 | 0.001*** | 193 | 0.001*** | 56.75 | 0.001 *** |
| Compartment (C) | 243 | 0.001*** | 55 | 0.001*** | 27.76 | 0.001 *** |
| Growth stage (S) | 57 | 0.001*** | 74 | 0.001*** | 18.66 | 0.001 *** |
| G:Y | 1.45 | 0.014* | 1.11 | 0.19 | 1.45 | 0.002 ** |
| G:C | 1.22 | 0.016* | 0.75 | 1.00 | 1.05 | 0.207 |
| G:S | 1.03 | 0.329 | 0.76 | 1.00 | 1.20 | 0.003 ** |
| Y:C | 51 | 0.001*** | 29.17 | 0.001 *** | 19.99 | 0.001 *** |
| Y:S | 18.27 | 0.001 *** | 47 | 0.001*** | 6.82 | 0.001 *** |
| C:S | 22.11 | 0.001 *** | 14 | 0.001*** | 6.94 | 0.001 *** |
| G:Y:C | 0.83 | 0.945 | 0.68 | 1.00 | 1.02 | 0.328 |
| G:Y:S | 0.82 | 0.88 | 0.63 | 1.00 | 1.03 | 0.333 |
| G:C:S | 0.70 | 1.000 | 0.51 | 1.00 | 0.84 | 1.000 |
| Y:C:S | 16.02 | 0.001*** | 12 | 0.001*** | 3.85 | 0.001 *** |
| G:Y:C:S | 0.80 | 0.98 | 0.62 | 1.00 | 0.98 | 0.523 |

‘\*\*\*’ : P<0.001, ‘\*\*’ : P<0.01 ‘\*’: P< 0.05 ‘.’: P<0.1.

The ordinations based on the fungal ITS region amplicon ASVs showed a slightly different picture, with an overwhelming effect of sampling year (2013-2014 vs 2015-2016) and growth stage (Fig. 1). Within the clusters due to sampling year, a clear effect of compartment could be observed, with root samples separated from rhizosphere and bulk soil samples (Fig. 1). Here again, this visual interpretation was confirmed by the Permanova analysis, where Year had the strongest effect followed by Growth stage, Compartment and then Genotype (Table 2). These main effect terms were all highly significant, except for Genotype that was marginally significant (Table 2). Some interaction terms were also significant, but none of them included Genotype.

The ordination based on the cpn60 gene amplicon ASVs showed a picture in between the ones observed for the 16S rRNA gene and ITS region amplicons (Fig. 1). Compartment and Year clearly had a strong effect on the microbial community structure, but these two terms seem to interact, with a more distinct root community structure for the 2013 samples (Fig. 1). The 2015-2016 showed here again a tight clustering, but where also joined by the 2014 samples (Fig. 1). In Permanova analyses, Year had the strongest effect, followed by Compartment, Growth stage and Genoptype, which were all highly significant (Table 2). Several interaction terms were also significant, notably Genotype:Year and Genotype:Growth stage (Table 2).

### Effect of genotype

Because of the significant Year x Compartment x Growth stage interaction for all three amplicon datasets groups and to explore more deeply our initial hypothesis about genotypes, Permanova analyses for the effect of genotype were performed separately for each Year x Compartment x Growth stage combination (Table 3). Since this reduced the number of samples in each analysis and consequently, the statistical power of our approach, we are also reporting and discussing test results that were 0.05<P<0.10. For bacterial communities (16S rRNA gene), the effect of genotype in the roots was significant at all growth stages for year 2013 and 2014, but only at dough development for 2015 and at stem elongation for 2016 (Table 3). The only occurrence where genotype was significant in the rhizosphere for 16S was at stem elongation in 2015 (Table 3). In contrast, the effect of genotype on fungal communities (ITS) was only significant for the 2013 samples, in the roots for all growth stages and in the rhizosphere for the dough development growth stage (Table 3). The patterns of significance for the cpn60 gene dataset were like the ones observed for the 16S rRNA gene dataset, with significant effects of genotype on root communities in 2013 (both growth stages), 2014 (dough development), 2015 (stem elongation) and 2016 (dough development) (Table 3). Additionally, the genotype significantly affected the rhizosphere microbial communities at the stem elongation stage in 2016 (Table 3).

**Table 3.** Permanova analyses for the effect of genotype for each year/compartment/growth stage combinations for 16S, ITS and cpn60 amplicon datasets generated from bulk soil, wheat rhizosphere soil and wheat roots collected from 2013-2016 at the stem elongation and dough development stages.

| Year |  | 16S |  | ITS |  | cpn60 |  |
| --- | --- | --- | --- | --- | --- | --- | --- |
| Compartiment | Growth stage | *F* | *P* | *F* | *P* | *F* | *P* |
| **2013** |  |  |  |  |  |  |  |
| Bulk | SE | 0.99 | 0.46 | 0.89 | 0.77 | 0.93 | 0.68 |
|  | DD | 1.04 | 0.32 | 0.86 | 0.90 | 1.00 | 0.47 |
| Rhizosphere | SE | 0.78 | 0.96 | 0.92 | 0.77 | 1.05 | 0.32 |
|  | DD | 1.05 | 0.35 | **1.25** | **0.06** | 1.03 | 0.36 |
| Roots | SE | **1.17** | **0.09** | **1.26** | **0.03\*** | **1.13** | **0.03\*** |
|  | DD | **1.16** | **0.09** | **1.41** | **0.03\*** | **1.12** | **0.08** |
| **2014** |  |  |  |  |  |  |  |
| Bulk | SE | 0.87 | 0.96 | 0.99 | 0.5 | 1.02 | 0.47 |
|  | DD | 0.94 | 0.76 | 0.87 | 0.94 | 0.97 | 0.57 |
| Rhizosphere | SE | 0.90 | 0.83 | 1.06 | 0.22 | 1.04 | 0.41 |
|  | DD | 0.98 | 0.52 | 1.06 | 0.29 | 0.98 | 0.53 |
| Roots | SE | **1.24** | **0.05** | 0.97 | 0.58 | 1.04 | 0.25 |
|  | DD | **1.24** | **0.01\*** | 0.94 | 0.72 | **1.18** | **0.01\*** |
| **2015** |  |  |  |  |  |  |  |
| Bulk | SE | 0.99 | 0.44 | 0.71 | 0.88 | na | na |
|  | DD | 0.86 | 0.80 | 0.71 | 0.82 | na | na |
| Rhizosphere | SE | **1.24** | **0.05** | 0.62 | 0.97 | 1.45 | 0.26 |
|  | DD | 1.04 | 0.34 | 0.52 | 0.99 | na | na |
| Roots | SE | 0.93 | 0.63 | 0.60 | 0.96 | **1.28** | **0.09** |
|  | DD | **1.33** | **0.05** | 0.85 | 0.76 | na | na |
| **2016** |  |  |  |  |  |  |  |
| Bulk | SE | na | na | 0.66 | 0.76 | na | na |
|  | DD | 1.01 | 0.44 | 0.44 | 0.99 | 1.08 | 0.25 |
| Rhizosphere | SE | na | na | 0.58 | 0.97 | **1.41** | **0.06** |
|  | DD | 0.93 | 0.62 | 0.56 | 0.98 | 1.16 | 0.15 |
| Roots | SE | **1.34** | **0.06** | 0.92 | 0.61 | 1.07 | 0.27 |
|  | DD | 1.15 | 0.14 | na | na | **1.15** | **0.09** |

‘\*\*\*\*’ : P<0.001, ‘\*\*\*’ : P<0.01 ‘\*’: P< 0.05 ‘.’: P<0.1.
SE: stem elongation; DD: dough development.

We further explored the effect of genotype on the phylum/class level community composition in the roots for the growth stages where genotype was significant in Permanova. For the 16S rRNA gene, ANOVA revealed that, in the roots, Actinobacteria (2013 dough development stage), Alphaproteobacteria (2015 dough development stage), Gammaproteobacteria (2013 and 2014 stem elongation stage and 2015 dough development stage) were significantly affected by genotype (Table S1 and Fig. 2). In Tukey HSD post-hoc tests, we observed that many of the significant differences observed were between *durum* and *aestivum* wheat genotypes (Table S1). For fungi, wheat genotypes significantly affected the relative abundance of Ascomycota and Basidiomycota in the roots at stem elongation in 2013 (Table S2 and Fig. 3), and the only significant difference in Tukey HSD post-hoc tests was between the relative abundance of Ascomycota between Marquis and CDC Verona (Table S2). Finally, for the cpn60 dataset, Acidobacteria (2013, 2014, 2016), Actinobacteria (2013), Alphaproteobacteria (2016), Bacteriodetes (2016) and Verrucomicrobia (2016) were significantly affected by the genotype in the roots at the dough development stage (Table S3 and Fig. 4). Tukey HSD post-hoc tests showed that some of the significant differences were between the older genotypes (Pelissier, Red Fife and Marquis) and some other more recent genotypes (Table S3).

**Figure 2.**
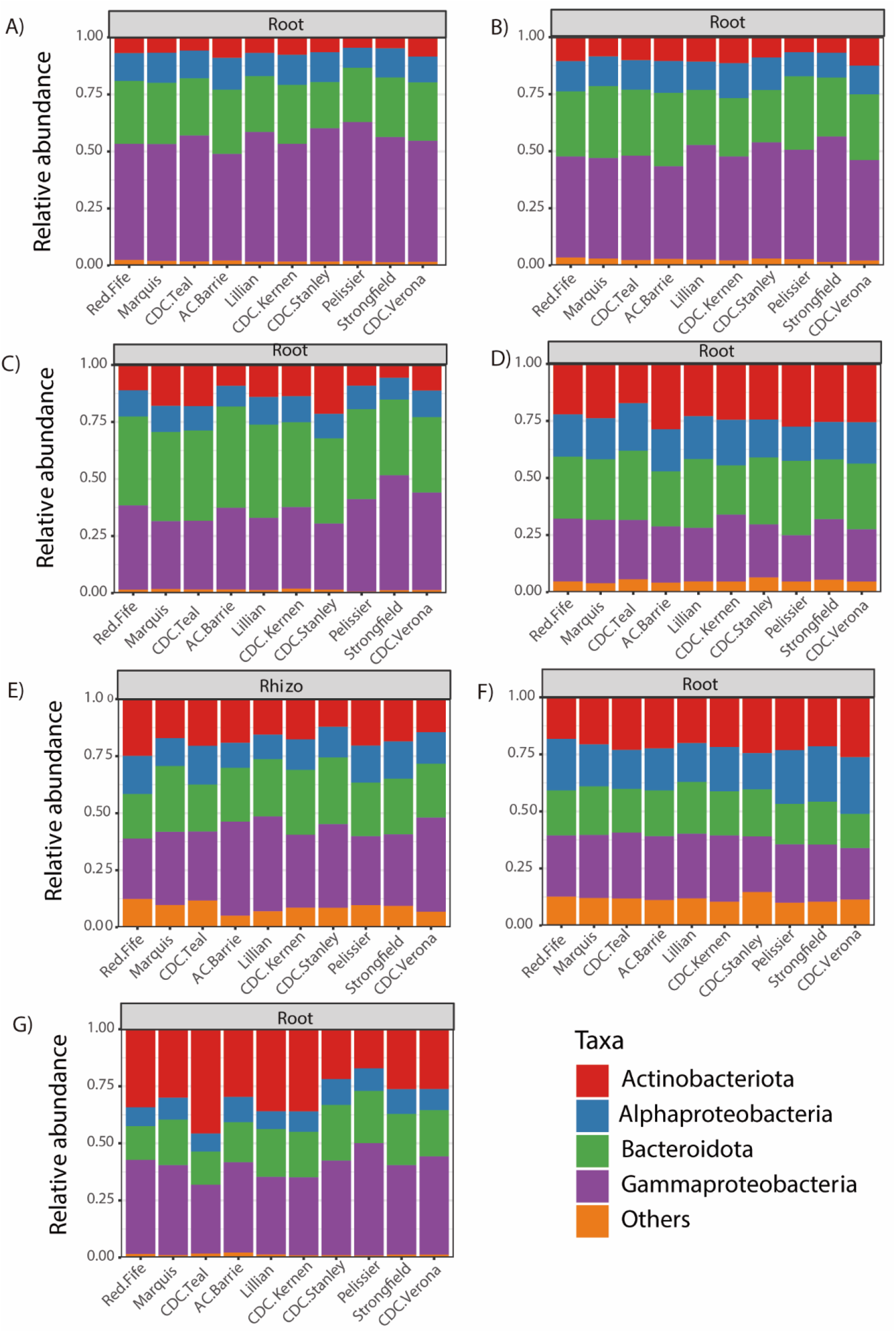
Community composition of the most abundant phylum/class based on the 16S rRNA gene amplicons for the growth stage/compartment combinations that showed a significant (P<0.10) effect of genotypes in Table 3. A) 2013, roots, stem elongation; B) 2013, roots, dough development; C) 2014, roots, stem elongation; D) 2014, roots, dough development; E) 2015, rhizosphere, stem elongation; F) 2015, roots, dough development; G) 2016, roots, stem elongation.

**Figure 3.**
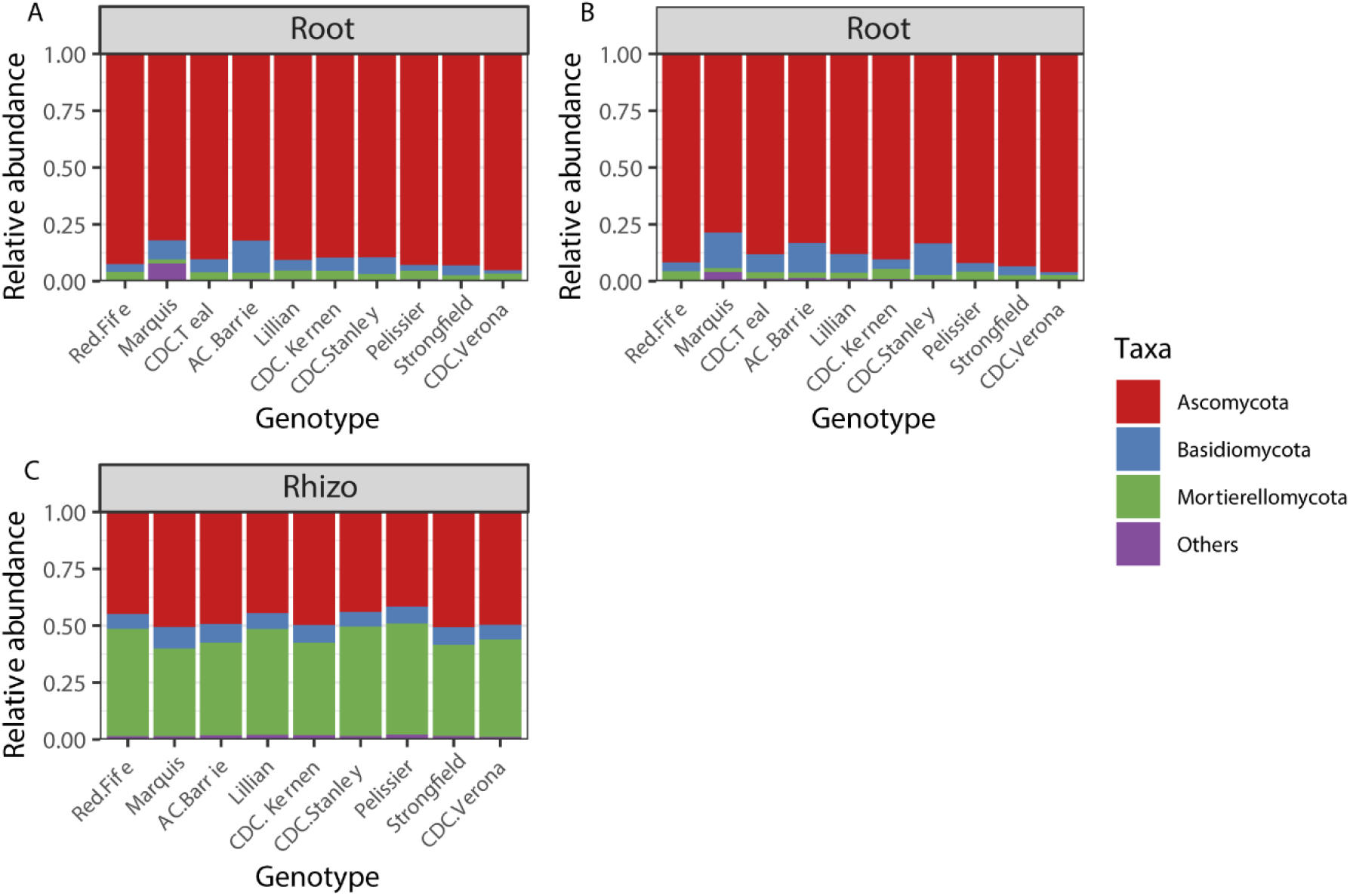
Community composition of the most abundant phylum based on the ITS region amplicons for the growth stage/compartment combinations that showed a significant (P<0.10) effect of genotypes in Table 3. A) 2013, roots, stem elongation; B) 2013, roots, dough development; C) 2013, rhizosphere, dough development.

**Figure 4.**
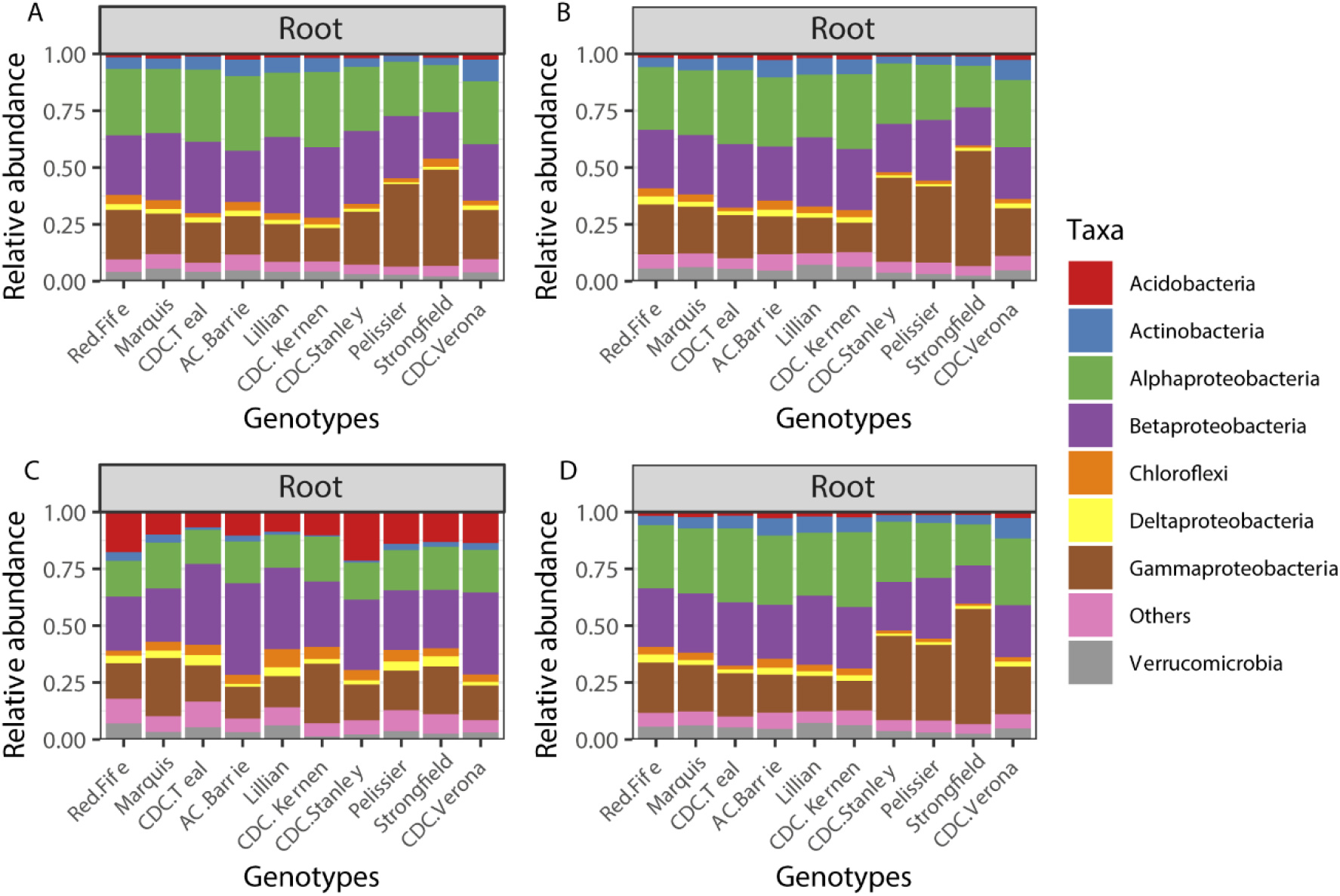
Community composition of the most abundant phylum/class based on the cpn60 gene amplicons for the growth stage/compartment combinations that showed a significant (P<0.10) effect of genotypes in Table 3. A) 2013, roots, stem elongation; B) 2013, roots, dough development; C) 2014, roots, dough development, D) 2016, roots, dough development.

## Discussion

Our multi-year study of the microbiome of different wheat genotype revealed an overwhelming influence of time, with sampling year and growth stage strongly influencing the bacterial and fungal communities associated with wheat root and rhizosphere. In addition, the microbiome of the roots was generally well differentiated from the one inhabiting the rhizosphere and bulk soil. Within these large temporal and spatial effects, we could still detect a significant effect of wheat genotype, which, in agreement with our hypothesis, varied through time and was mostly significant for the root microbiome. We had previously constructed metagenomic libraries from the rhizosphere samples collected from this experiment at the stem elongation stage in 2013. The lack of effect from the wheat genotype on the metagenome presented in Quiza et al. [9] is highly coherent with the results presented here, as no significant effect of genotype could be observed for 16S, ITS nor cpn60 in the rhizosphere samples collected at the stem elongation stage in 2013. However, when expanding our sampling, we realized that, for genotype, significance was mostly seen for roots samples, and even then, it varied through the years and wheat growth stages. These results reconcile Quiza et al. [9] with the literature that reported significant effects of genotype for wheat microbial community structure, composition and functions [19, 40–42], by suggesting that the samples taken for the Quiza et al [9] study were not affected, but that at other times or in different compartments, this effect would have been apparent. Alternatively, it could very well be that, because of the high functional redundancy of microbial communities, the changes observed here at the taxonomic level have little effect on the functional makeup of the community and would have been undetectable using shotgun metagenomics.

Here, we report for the first time that the effect of genotype on the wheat microbiome in the field is highly variable in both space and time, with more significant effects in the root compartment. This variability in the genotype-specific effects on root-associated microbial communities challenges the view that modern genotypes would have lost their ability to associate with nutrition-beneficial microbial communities when grown under low inputs [4–8]. Indeed, for root- and rhizosphere-associated bacteria and fungi of wheat grown under low nutrient conditions, most of the sample categories did not show an influence of genotype, suggesting that if this loss in the ability to associate with beneficial microorganisms could potentially happen, it is highly transient. Although changes in function could have potentially happened without concomitant changes in marker gene data, our data is highly coherent with the shotgun metagenomic study previously carried out on a subset of our samples [9]. Alternatively, the relatively short genotype development gradient (~175 years) might have precluded the observation of more significant trends. However, previous studies that found a clear genotype effect were mostly done using controlled growth conditions, or on a single sampling date or single compartment, and it is thus difficult to conclude if our observations would also apply to other crops or along a longer genotype development gradient (e.g. by including wild parental plants).

In sharp contrast with many previous reports, where fungi are often reported to be more intimately linked to plants, and therefore more influenced by variations between genotypes [51] whereas bacteria are more often primarily influenced by soil properties [40, 52, 53], we found here that bacteria were more strongly influenced by wheat genotype than fungi. In fact, in general permanova tests, there was no main or interactive effect of genotype on fungal communities, and in more in-depth anova analyses, we only found an effect of genotype on the fungal communities of the root and rhizosphere of wheat in 2013. Another evidence of the less intimate linkage between fungi and the plant is the fact that fungal communities were overwhelmingly influenced by sampling year, and very little by compartment, whereas bacteria were more strongly influenced by plant compartment than by sampling year. It was shown that the influence of genotypes on fungal communities increases with stress [40, 51], suggesting that the low nutrient conditions under which wheat was planted in the current field experiment were not stressful. However, Quiza et al. [9] reported a decrease in yields as compared to expected yields for the different genotypes under high input conditions, which would indicate some nutrient limitation.

Plant exudation patterns vary with plant development stages [17], which results in shifts in microbial communities and activities [11, 12]. Although these previous studies were performed under strictly controlled experimental conditions, plant development stage could also have played a role in the seasonal patterns observed here, although it is difficult to disentangle from the influence of fluctuating environmental conditions under field conditions. Indeed, the changes observed with wheat growth stage could be due to the direct and indirect (through plant) influence of changing environmental conditions on microbial communities. These fluctuations in environmental conditions could also explain the year-to-year variations observed in microbial communities. In fact, fungal and bacterial communities clustered together based on sampling year (2013-2014 vs 2015-2016), and this appear to have changed the effect of genotype which was more often significant in 2013-2014 than in 2015-2016 (10 out of the 16 significant occurrences vs 6 out of 16, respectively). When comparing historical weather data, the month of June was wet and cold for both 2013 and 2014 (117.7 and 94.8 mm of rain, average daily temperature of 20.9°C and 19.4°C, respectively), as compared to 2015 and 2016 (13.6 and 49.7 mm of rain, average daily temperature of 24.8°C and 24.7°C, respectively) (Table 1). Conversely, the rest of the summer was wetter in 2015 and 2016 (179.5 mm and 152.9 mm of rain, respectively) than in 2013 and 2014 (65.9 mm and 73.7 mm of rain, respectively). Precipitation is often cited as an important factor shaping soil microbial communities, in view of its influence on soil water content, gas diffusion and redox conditions and the soil processes that they influence. Recently, we showed that microbial communities, and most especially archaeal ammonia-oxidizers, shifted dramatically following a large drying-rewetting event, whereas little change was observed in response to small changes in average soil water content [19]. Soil water stress history is also a major influencing factor for wheat microbial communities [41].

In our study, the effect of genotype did not extend past the rhizosphere, as in no case was there a genotype effect recorded in bulk soil samples. Recent studies had highlighted that an effect of genotype could be seen in the bulk soil [19, 40, 42], which led to the suggestion that because of the effect of gaseous compounds, the rhizosphere could in practice extend past the few mm of soil surrounding the roots [54]. Alternatively, fungal hyphae extending from the rhizosphere could also have played a role in these previous studies. However, here, as expected, the effect of genotype was more often significant for the root communities, as 13 out of the 16 significant effects of genotype were observed for root communities. Previous studies showed that genotype effect of wheat increased in the following order rhizosphere<roots<leaves [40], which makes sense as the strength of the plant selective pressure is likely to increase inside plant tissues.

Overall, although we did find some significant influence of genotype on root and rhizosphere microbial communities, this effect varied with time and was mostly visible for bacteria and for root samples. This suggests that the development of modern wheat cultivars did not result in stable shifts in the root-associated microbial communities, and that these changes are anyway dwarfed by the shifts caused by changing environmental conditions and plant development stages that occur throughout and across the years. This knowledge is highly relevant for microbiome engineering approaches [13, 55], as it highlights the overwhelming strength of environmental selection vs. plant selection.

## Acknowledgements

This work was supported by the Wheat Flagship Program of the National Research Council Canada and by Natural Sciences and Engineering Research Council Discovery grants to EY (RGPIN-2014-05274) and to MSA (RGPIN-2014-05426). We would like to thank Sylvie Sanschagrin for technical help in the laboratory. We acknowledge Compute Canada for access to the Guillemin (McGill University) and Graham (University of Waterloo) systems.

---

**Supplementary Table S1.**
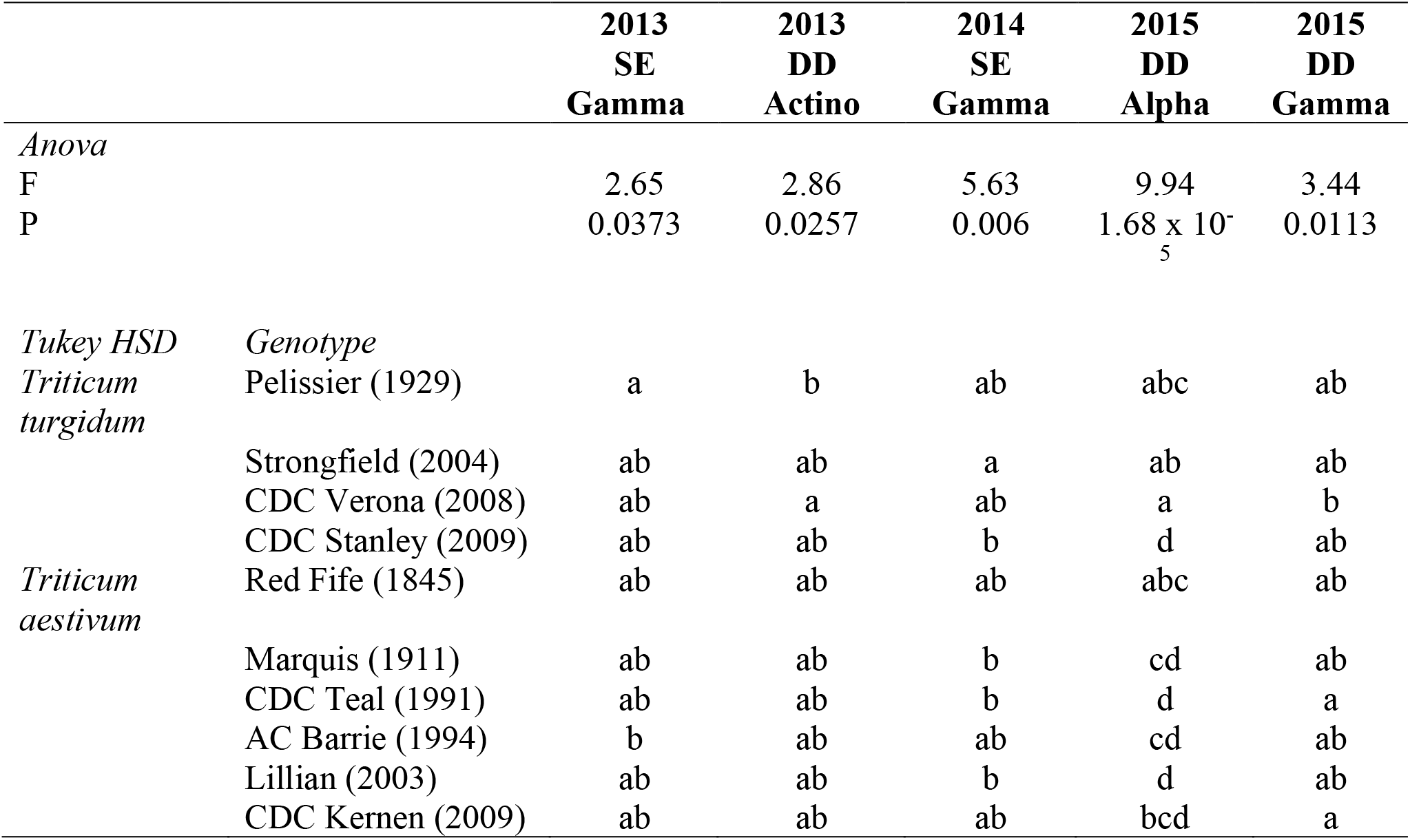
Anova tests and Tukey HSD post-hoc tests for the effect of genotype on the relative abundance of bacterial phylum/classes in the roots based on the 16S rRNA gene amplicon dataset.

**Supplementary Table S2.**
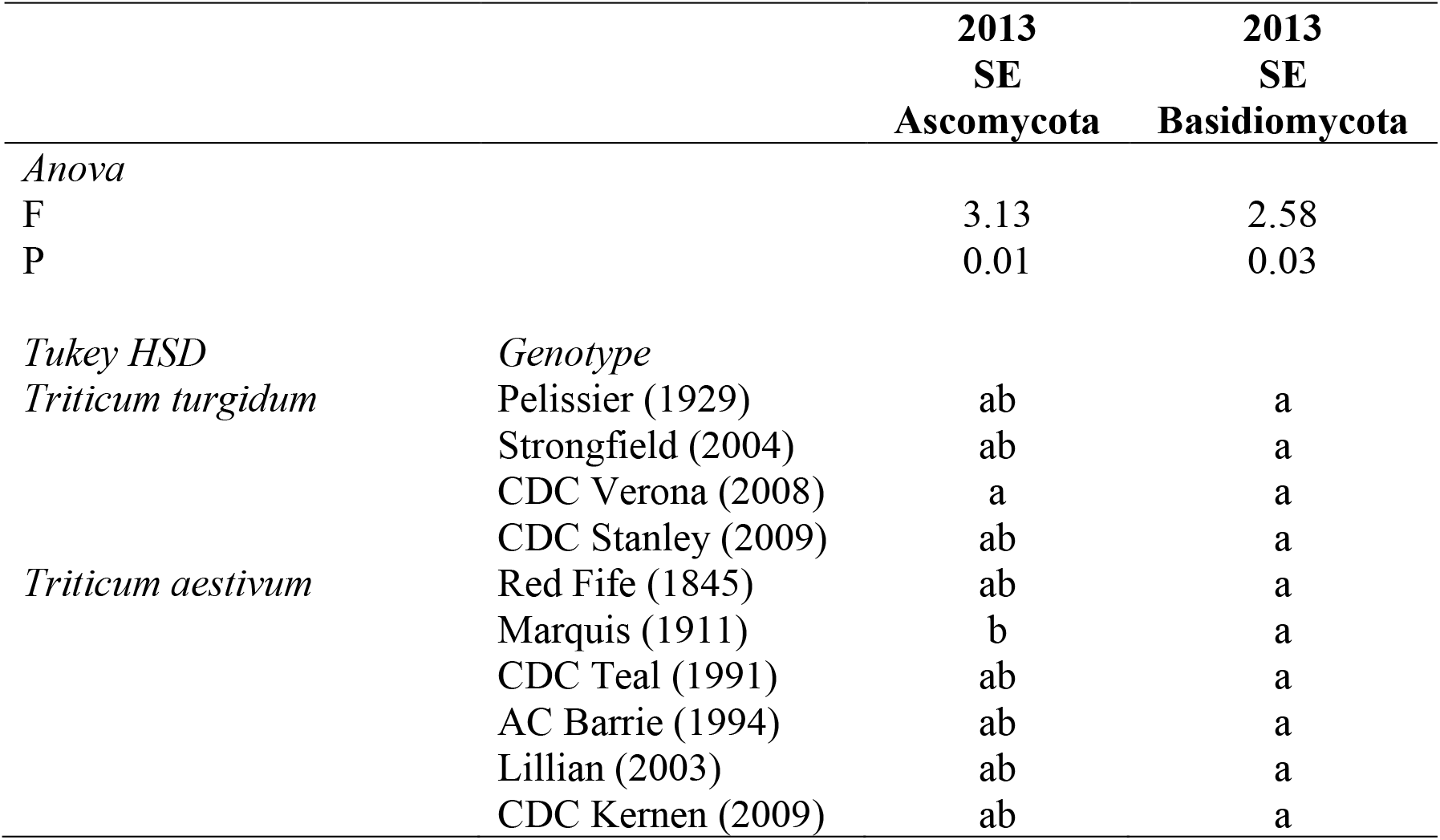
Anova tests and Tukey HSD post-hoc tests for the effect of genotype on the relative abundance of fungal phyla in the roots based on the ITS region amplicon dataset.

**Supplementary Table S3.** Anova tests and Tukey HSD post-hoc tests for the effect of genotype on the relative abundance of bacterial phylum/classes in the roots based on the cpn60 gene amplicon dataset.

|  |  | 2013<br>DD<br>Acido | 2013<br>DD<br>Actino | 2014<br>DD<br>Acido | 2016<br>DD<br>Acido | 2016<br>DD<br>Alpha | 2016<br>DD<br>Bacteroid | 2016<br>DD<br>Verruc |
| --- | --- | --- | --- | --- | --- | --- | --- | --- |
| *Anova* |  |  |  |  |  |  |  |  |
| F |  | 2.51 | 3.15 | 2.47 | 4.59 | 3.44 | 4.75 | 2.79 |
| P |  | 0.0487 | 0.0199 | 0.0489 | 0.006 | 0.0217 | 0.005 | 0.045 |
| *Tukey HSD* |  |  |  |  |  |  |  |  |
| *Triticum turgidum* | *Genotype*<br>Pelissier (1929) | a | b | ab | ab | ab | b | a |
|  | Strongfield (2004) | a | ab | ab | ab | ab | b | a |
|  | CDC Verona (2008) | a | a | ab | ab | ab | ab | a |
|  | CDC Stanley (2009) | a | ab | a | ab | ab | b | a |
| *Triticum aestivum* | Red Fife (1845) | a | ab | ab | ab | a | b | a |
|  | Marquis (1911) | a | ab | ab | ab | ab | a | a |
|  | CDC Teal (1991) | a | ab | b | a | ab | b | a |
|  | AC Barrie (1994) | a | ab | ab | ab | ab | b | a |
|  | Lillian (2003) | a | ab | ab | a | b | b | a |
|  | CDC Kernen (2009) | a | ab | ab | ab | b | b | a |

